# Adopting orphan receptors: group IX/X transition metals as potential ligands of zebrafish Tlr4 homologs

**DOI:** 10.1101/2023.09.13.557392

**Authors:** Aaron P.D. Fox, Tracy S. Lee, Sakina N. Mithaiwala, Niall M. Pollock, Amit P. Bhavsar, W. Ted Allison

## Abstract

TLR4 is the prototype immune receptor and central to infection defence via detecting lipopolysaccharide (LPS). Surprisingly, the impacts of LPS upon the TLR4 homologs in zebrafish, an important animal model, are equivocal and the function of TLR4 homologs across all fishes remains debatable. Recent work suggests zebrafish Tlr4 mediates ototoxic responses to a platinum-based chemotherapeutic. This prompts our hypothesis that Tlr4 detects group IX/X transition metals and thus has conserved roles with human TLR4 mediating allergic responses to nickel. Here, we use the larval zebrafish lateral line model to demonstrate (sub-)micromolar Ni, Co and Pt are ototoxic in a dose-dependent manner. TLR4 homologs are required for this toxicity because Tlr4 knockdown via CRISPR significantly reduced the metals’ impacts by ∼50%. Moreover, zebrafish Tlr4 was sufficient to mediate inflammatory responses to metals when expressed in a human cell line. These data are consistent with the notion that mediating responses to transition metals was a function of TLR4 homologs in the last common ancestor of fish and mammals, and begins to resolve the function(s) of TLR4 homologs in the zebrafish animal model of disease.

## 1. Introduction

Toll-like receptors (TLRs) are evolutionarily ancient proteins with origins dating back to more than 700 MYA and can be found in a range of organisms from corals to humans (Behzadi et al., 2021; Fitzgerald & Kagan, 2020). Mammalian TLR4 is multifaceted in its function with involvement in pathogen recognition, cancer pathology, and autoimmune disease (Heine & Zamyatina, 2022). The best studied function of TLR4 in humans is its ability to detect and bind to microbial pathogen associated molecular patterns (PAMPs) such as lipopolysaccharide (LPS), and viral glycoproteins, which are essential for the body’s inflammatory response to pathogens. The structure of TLR4 is conserved between species, however the function of TLR4 homologs in zebrafish (and in all fish and other early-branching vertebrates) has remained elusive. This has potential to frustrate interpretations when using zebrafish as an important model of disease, and so we seek to fill that knowledge gap here.

TLR4 proteins can be classified as pattern recognition receptors (PRRs), and like all PRRs they act as the sentinels of the innate immune system, surveying the host environment for signs of foreign molecules and alarming the host to damage or foreign bodies. Mammalian TLR4’s primary ligand is LPS, an important extracellular component of the cell wall in Gram-negative bacteria. In zebrafish, however, no convincing evidence demonstrates that LPS signals via TLR4; although LPS has impacts when applied to zebrafish (often only at extra-biologic high concentrations, and in screening efforts to discover phytobiologicals) as demonstrated in scores of papers (Athapaththu et al., 2022; Bates et al., 2007; Fernando et al., 2017; Hwang et al., 2016; Lee et al., 2013; Li et al., 2022; Urzi et al., 2023; Yang et al., 2014; Yu et al., 2022; Zhang et al., 2018), these impacts of LPS have not been shown to require TLR4. The reason for the disparity between TLR4 signalling in mammals and fish may be attributed to TLR4 itself and/or to the evolution of the associated intracellular signalling pathways (Candel et al., 2015; Candel et al., 2016; Liu et al., 2010; Martinez-Lopez et al., 2023; Novoa et al., 2009; Palti, 2011), or perhaps the appropriate source of LPS has yet to be tested. In mammals, signalling through TLR4 using LPS requires multiple different co-receptors acting in concert with one another including: LPS binding protein (LBP), cluster differentiation 14 (CD14), and myeloid differentiation factor 2 (MD-2) (Ryu et al., 2017). Activation of TLR4 leads to downstream signalling and initiation of pro-inflammatory and type 1 interferon related gene expression through NF-κB and IRF3 respectively (Figure 1). Dysfunction in any of these signalling components has been demonstrated to impede the proper functioning of TLR4 as a bacterial sensor (Ciesielska et al., 2021). Recently, the homolog for MD-2 was discovered in zebrafish, prompting Loes et al. to reconsider Tlr4’s role in LPS recognition and signalling within zebrafish (2021). Although Loes et al. found the zebrafish homolog Tlr4ba was capable of being activated by high concentrations of LPS, it’s low sensitivity to LPS and the lack of other crucial co-receptors in the zebrafish genome suggests zebrafish Tlr4 primarily recognizes other unidentified ligands. MD-2 contributes to signalling through LPS, but it also plays a role in signalling through other TLR4 ligands such as metals. Overall, the detection of LPS via TLR4 appears to have evolved in mammals after that lineage split from other vertebrates, and thus the function of TLR4 in fishes, including those related to important models of disease, has remained elusive (Martinez-Lopez et al., 2023).

**Figure 1.**
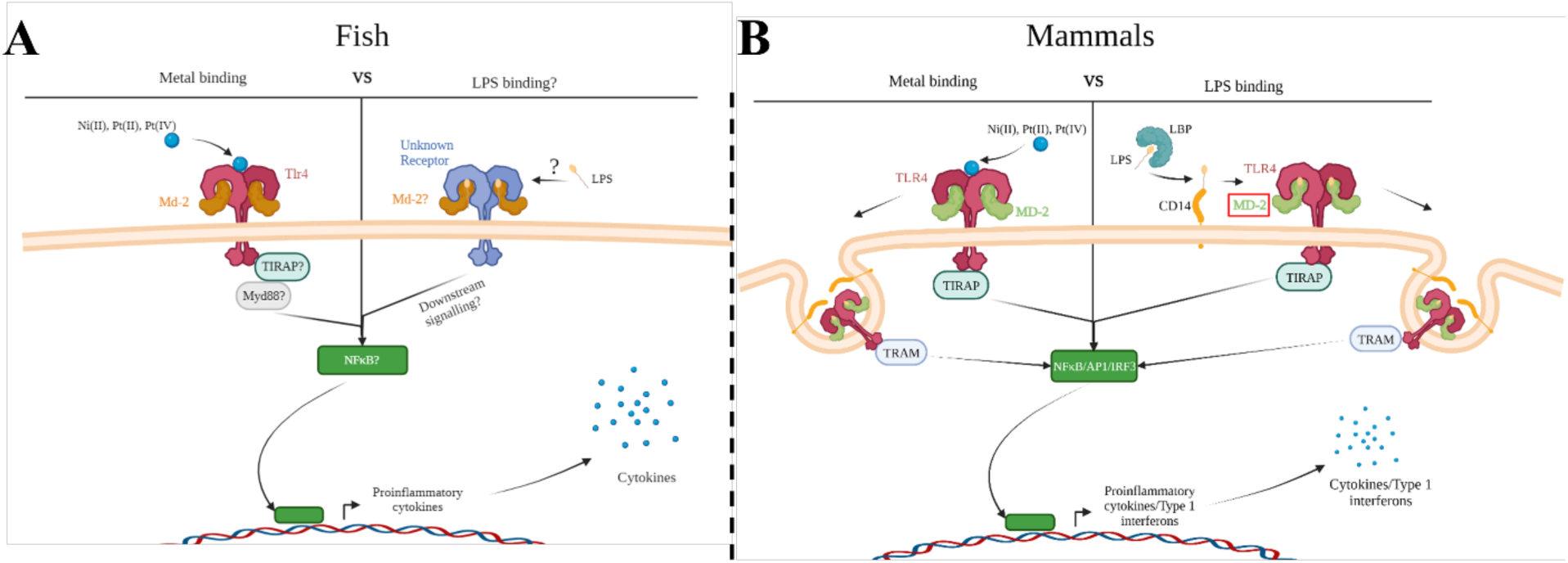
Metal ion activation of zebrafish Tlr4 and direct binding to human TLR4 versus co-receptor binding with LPS. Group IX/X transition metals activate a proinflammatory response through zebrafish toll-like receptor 4 (Tlr4) (A) or human TLR4 (B). **A**) Metal ions activate a proinflammatory response through zebrafish TLR4 homologs, which may be mediated through myeloid differentiation factor 2 (Md-2). Activation would likely lead to signalling through downstream molecules, such as myeloid differentiation primary response 88 (Myd88), leading to activation of a proinflammatory transcription factor (such as nuclear factor-κβ (NF-κβ)) and release of proinflammatory cytokines. LPS signalling in zebrafish remains unclear but would likely involve a similar signalling cascade as described above. **B**) Metal ions bind directly to key residues in the ectodomain of human TLR4 leading to dimerization and binding of MD-2. This stimulates nuclear factor-κβ (NF-κβ) and interferon regulatory transcription factor 3 (IRF3), leading to the release of type 1 interferons and proinflammatory cytokines. LPS binds to the LPS Binding Protein (LBP) and is transferred to the co-receptor cluster of differentiation 14 (CD14). CD14 facilitates the binding of LPS to MD-2/TLR4 signalling complex, which then dimerizes and follows a similar signalling cascade as described above. Created with BioRender.

The observation that metals, such as nickel, can act as ligands of TLR4 was a crucial discovery in determining the involvement of the innate immune system in mediating metal contact hypersensitivity, a type of allergic contact dermatitis (ACD) (Schmidt et al., 2010). Two mechanisms are believed to play a role during the sensitization of the immune system to nickel (Riedel et al., 2021). One mechanism involves the generation of an adaptive immune response following first contact with the allergen, while the other mechanism is through the activation of innate immune cells via TLR4 (De Graaf et al., 2023; Riedel et al., 2021; Saito et al., 2016). Similar to LPS, metal binding to TLR4 leads to the production of pro-inflammatory cytokines, recruiting immune cells and generating inflammation resulting in itching, tenderness, swelling, and rashes (Katsnelson, 2010; Saito et al., 2016). However, metal binding to TLR4 is through a distinct mechanism that doesn’t require the co-receptors associated with LPS signalling (Domingo et al., 2023; Peana et al., 2017; Raghavan et al., 2012) (Figure 1). Metals bind directly to the ectodomain of TLR4 leading to TLR4 dimerization through a cluster of conserved histidine residues (Oblak et al., 2015; Peana et al., 2017). Oblak et al. determined MD-2 was only needed for stabilization of the TLR4 complex during metal ion binding after dimerization and allows for downstream signalling through the MyD88-dependent and -independent pathways (2015). Taken together, we were motivated to investigate metals as potential ligands of zebrafish Tlr4 homologs (Loes et al., 2021).

Zebrafish are a potent model organism for the study of human disease and have gained recognition as an excellent model for immunological processes (Gomes & Mostowy, 2020; van der Sar et al., 2004) and drug discovery (Patton et al., 2021; Stewart et al., 2014; Yang et al., 2014). They allow for examination of cellular interaction at a whole-organism level and possess many of the same innate immune components as humans (Gomes & Mostowy, 2020; Novoa & Figueras, 2012). Zebrafish possess a total of 20 TLRs and express three TLR4 homologs known as Tlr4ba, Tlr4bb, and Tlr4al. Unlike mammalian TLR4, zebrafish Tlr4 has not been demonstrated to respond to LPS, and zebrafish lack the co-receptor CD14. LPS is a modular structure that can be capped by an O-antigen, referred to as “smooth” LPS, where CD14 is required for TLR4 activation by smooth LPS in mammals (Martinez-Lopez et al., 2023; Sepulcre et al., 2009; Sullivan et al., 2009). Nevertheless, zebrafish TLR4 proteins are structurally congruent with their mammalian counterparts, with all three homologs coding for an extracellular leucine-rich repeat (LRR) ectodomain, a transmembrane domain and a toll/interleukin-1 receptor (TIR) containing domain (Vaure & Liu, 2014) (Supplementary Figure 1). Moreover, Loes et al. has recently uncovered the existence of a zebrafish Md-2 homolog that interacts with Tlr4ba, providing evidence of an ancestral interaction between Tlr4 and Md-2 facilitating the detection of extracellular ligands (2021). In chimeric TLR4 fusion experiments, their transmembrane and TIR domains have also shown to function in a similar manner to mammals, capable of eliciting a proinflammatory response after activation (Sullivan et al., 2009). Furthermore, Purcell et al. showed zebrafish and humans possess many of the same intracellular signalling molecules such as MyD88, TIRAP, TRIF, TRAF6, IRF3, and IRF7 (2006). The difference in their ligand specificity appears to lie within the ectodomain, with Tlr4ba, Tlr4al, and Tlr4bb proteins showing ∼36% identity to the ectodomain of human TLR4 respectively (Supplementary Figure 2).

The lack of a known primary ligand for zebrafish Tlr4 is puzzling as they appear to have all the components required for sensing an extracellular ligand, yet their true function is still a mystery. Our previous studies have shown zebrafish Tlr4 to be involved in cisplatin-induced ototoxicity (Babolmorad et al., 2021). Morpholino knockdown of *tlr4ba* and *tlr4bb* mitigated cell death induced by the platinum (II) based chemotherapeutic cisplatin (Babolmorad et al., 2021). This observation, and the unique mechanism of TLR4 activation by metals in mammals, led us to hypothesize that group IX/X transition metals are a ligand of TLR4 that predates the evolutionary split between fish and mammals. Thus, we predict that group IX/X transition metals will initiate zebrafish Tlr4 activation, leading to NFκβ induction and a proinflammatory response. Here, we present Ni (II), Co (II), Pt (II) and Pt (IV) as novel activators and potential ligands of zebrafish Toll-like receptor 4 (Tlr4). The findings improve the understanding of Tlr4 evolution, and ultimately advance the utility of zebrafish as an animal model for future studies in innate immunity and disease.

## 2. Methods

### 2.1 Zebrafish Husbandry and Ethics

Zebrafish were kept in standard conditions at the University of Alberta aquatics facility following a 14:10 light/dark cycle in 28.5°C. They were fed twice daily using either trout chow or brine shrimp. All zebrafish were raised, bred, and maintained following the institutional Animal Care and Use Committee approved protocol AUP00000077 which operates under guidelines set by the Canadian Council of Animal Care.

### 2.2 Zebrafish breeding and care

Wildtype (AB strain) zebrafish were bred and maintained at 28.5°C in standard conditions (Westerfield, 2007). Embryos were maintained in E3 media containing 0.01% methylene blueE3 media was made from a 60X stock solution of embryo media (prepared with 0.29M NaCl, 0.01M KCl, 0.026M CaCl_2_, 0.001M MgSO_4_• 7H_2_O) diluted to 1X in Milli-Q water or mixed with 0.01% methylene blue solution in a 5.5:1 ratio before being diluted in Milli-Q water. Embryos were grown in a 28.5°C incubator and the E3 media was replaced daily.

### 2.3 Metal ion treatment of larval zebrafish

Wildtype (AB strain) zebrafish larvae were grown to either 5- or 6-days post fertilization (dpf) in E3 embryo media with methylene blue and 10-15 larvae were placed in each well of a six-well plate. The larvae were then bath treated with either 0, 2.5, 5, 7.5, 10, or 15µM of nickel (II) (Sigma; prod. #654507) or cobalt (II) (Sigma; prod. #255599) chloride hexahydrate or 0, 0.25, 0.5, 0.75, 1, or 1.5µM of platinum (II) (Sigma; prod. #206091) or platinum (IV) (Sigma; prod. #206113) chloride diluted in E3 media with methylene blue for 20 hours at 28°C. All wells were then washed with embryo media three times before being incubated on an orbital incubator shaker at 130 rpm and 28°C for 15-20 minutes in media containing 0.01% 2-[4-(dimethylamino) styryl]-1-ethylpyridinium iodide (DASPEI, Sigma Aldrich; cat. #3785-01-1) to selectively stain live neuromast hair cells. The wells were washed twice using E3 media and groups of larvae were transferred into individual petri dishes where they were anaesthetized using 4% Tricaine-S (MS-222). After blinding the researcher to treatments, the neuromasts were imaged using a Leica M165 FC dissecting microscope equipped with a GFP-long pass fluorescent filter.

### 2.4 Neuromast quantification assay

The DASPEI quantification assay has been established by many studies, but the one used here was modified from Babolmorad et al. (2021; Coffin et al., 2009; Harris et al., 2003; Owens et al., 2007; Uribe et al., 2018). Briefly, five neuromasts along the posterior lateral line were chosen for scoring consistently throughout experiments (Figure 2). Each neuromast was assigned a score ranging from 0-2 based on fluorescent intensity. A score of 2 is assigned to bright neuromasts with no noticeable decline in fluorescence, 1.5 for minor decline, 1 for moderate decline, 0.5 for severe decline and 0 for complete loss of fluorescence. The neuromast viability score for each larva was determined by calculating the sum of these five scores, with a total maximum score of 10. A lower score represents a decline in neuromasts health (indicative of ototoxicity), while a higher score (near 10) represents healthy neuromasts.

**Figure 2.**
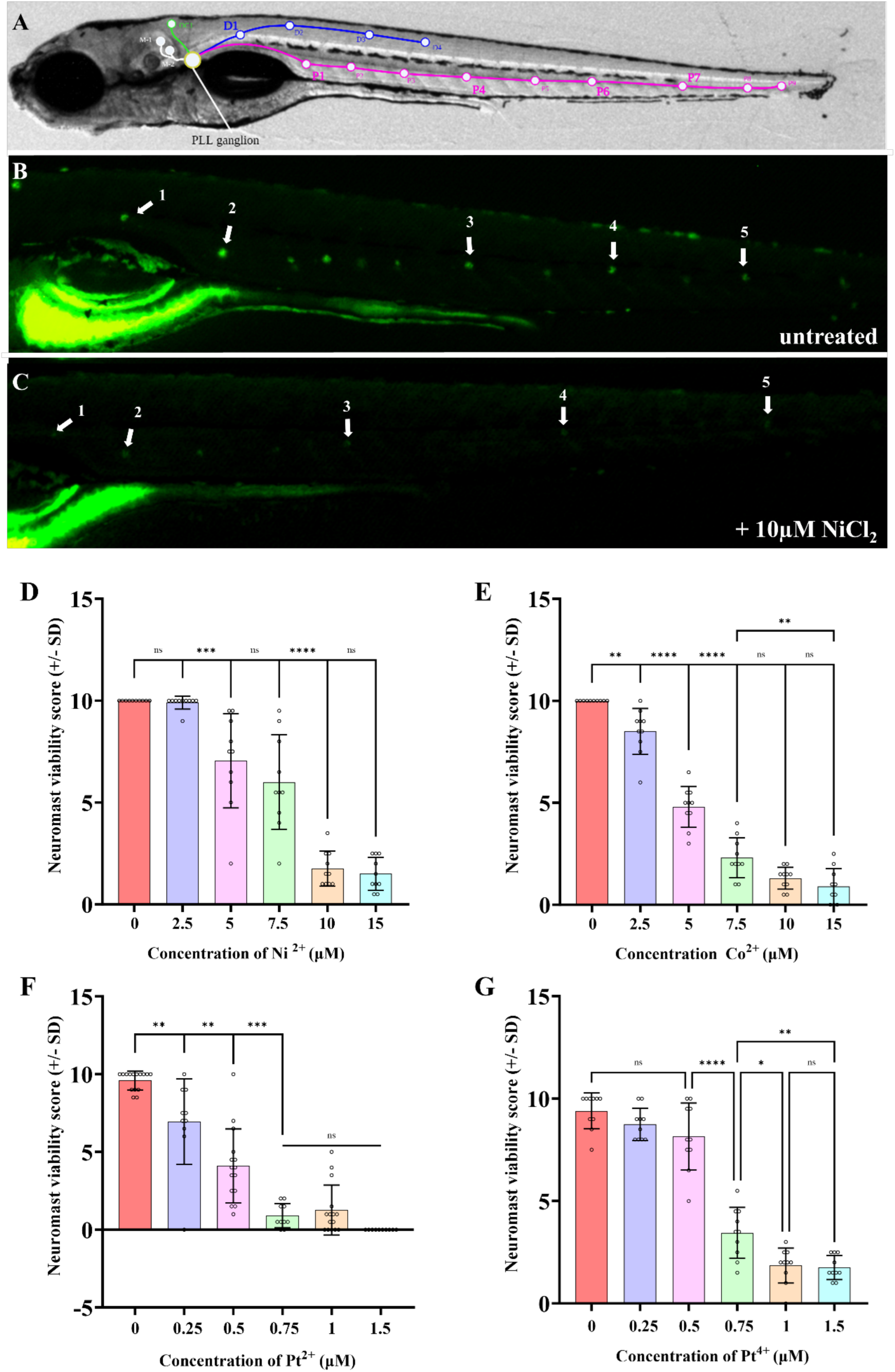
Nickel (II) chloride, cobalt (II) chloride, platinum (II) chloride, and platinum (IV) chloride induce PLL hair cell death in a dose dependent manner within 6-7dpf zebrafish. A) Location of posterior lateral line neuromasts drawn on a bright field image of a 7dpf larva. B) DASPEI stained untreated larva with visible neuromasts. White arrows indicate the neuromasts that were consistently selected for scoring between experiments. C) DASPEI stained larva treated with 10µM nickel (II) chloride hexahydrate, showing the significant decline in neuromast fluorescence due to hair cell death. White arrows indicate the selected neuromasts relative to those selected in panel A. As the concentration of the NiCl_2_ (D), CoCl_2_ (E), PtCl_2_ (F), and PtCl_4_ (G) in the media increased, the neuromast hair cell viability decreased. NiCl_2_ and CoCl_2_ were less toxic to neuromast hair cells compared to platinum salts. N = 10 wildtype larvae for all treatment groups. Bars represent the mean neuromast viability score of all larvae, and each dot represents data from an individual larva. ****p<0.0001, ***p<0.001, **p<0.01, *p<0.05, ns = not significant by one way ANOVA and Tukey’s multiple comparison test.

### 2.5 Guide RNA design and injections

Previous work has shown that the biallelic editing ability of clustered regularly interspaced short palindromic repeats (CRISPR)-cas9 technology allows for the generation of F0 “crispant” knockdowns in a target gene with high fidelity (>80%) (Burger et al., 2016; Hoshijima et al., 2019; Jao et al., 2013; Kroll et al., 2021; Shah et al., 2015). This approach allows for high throughput examination of loss-of-function phenotypes within zebrafish larvae with reduced number of resources and time. Knockdown of TLR4 homologs (the generation of *tlr4* “crispants”), used duplexes of trans-activating CRISPR-RNA (tracrRNA) and CRISPR-RNA (crRNA) to produce a single guide RNA (sgRNA). The cas9 nuclease binds to the tracrRNA and is guided to mutate the target gene through the crRNA. crRNAs for *tlr4ba*, *tlr4al,* and *tlr4bb* were designed using the predesigned Alt-R^®^ CRISPR-Cas9 guide RNA program offered by Integrated DNA Technologies (IDT). The crRNA sequences used can be found in Supplementary Table 1. The final gRNAs for each gene were made by mixing 100µM of the tracrRNA and 100µM of the crRNA, heating them to 95°C for 5 minutes, and diluting them to 25µM using the nuclease-free duplex buffer (IDT; cat. #11-05-01-12). All three gRNAs (25µM) were combined along with Cas9 protein (25µM) and heated to 37°C for 10 minutes to make a ribonuclease complex (25µM, RNP). The injection solution contained RNP (25µM), 1% dextran tetramethylrhodamine (invitrogen; cat. #D1817), 1M KCl, and RNAse-free water filled to 10µL. Wildtype (AB) zebrafish embryos at the 1 cell stage were injected with either a mock solution containing only the Cas-9 protein (mock injected) or the injection solution containing gRNAs (*tlr4ba/al/bb* crispant). Injected embryos were then incubated at 28.5°C in E3 media.

At 1 dpf dead embryos were removed from the petri dish and E3 media was replaced with fresh media. At 2 dpf embryos were screened for the injection using a Leica M165 FC dissecting microscope equipped with an mCherry fluorescent filter. Injected embryos showing red fluorescence were sorted and grown to 5 dpf at 28.5°C.

### 2.6 Confirming efficacy of tlr4 CRISPR via genotyping

A standard genomic DNA extraction was done on all uninjected, mock and *tlr4ba/al/bb* CRISPR injected larvae that were scored during metal experiments. The extraction was performed by submerging whole larvae in 15µL of 50µM NaOH and boiling them in the thermocycler at 95°C for 15 minutes, followed by a cooling period of 4°C for 5 minutes. 5µL Tris-HCl was added to neutralize the solution and DNA was stored at 4°C. A PCR was performed on genomic DNA samples using the AllTaq^TM^ Master Mix Kit (Qiagen; cat. #203146) following the manufacturer’s protocol. PCR product concentration was then quantified using a Nanodrop 2000 spectrophotometer (Thermo Scientific). Primers used for the PCR amplification of each zebrafish *tlr4* homolog can be found in Supplementary Table 1. Thermocycling conditions were a 2 minute denaturing period at 95°C, then a specific amplification period for five cycles of 94°C for 15 seconds, 64°C for 15 seconds, and 72°C for 30 seconds, followed by a less-specific amplification for 30 cycles of 94°C for 15 seconds, 54°C for 15 seconds, and 72°C for 30 seconds, and a final extension at 72°C for 10 minutes. Finally, Sanger-sequencing using the same PCR primer sets was performed on the amplified *tlr4ba*, *tlr4al*, and *tlr4bb* genes from 1 uninjected, 1 mock injected and 5 *tlr4* mutant larvae chosen at random.

### 2.7 HEK293T cell transfection and treatment

Human embryonic kidney (HEK) 293T cells (ATCC, catalog (cat) number CRL-3216) were grown in Dulbecco’s Modified Eagle Medium (DMEM) supplemented with FBS (10%) and penicillin-streptomycin (100 μg/mL) at 37°C and 5% CO_2_. Cells were seeded in 24-well plates (5 x 10^4^ cells/well) for functional assays and 6-well plates (5 x 10^5^ cells/well) for immunoblotting. 24 hours after seeding, the HEK293T cells were transfected with an empty vector (pcDNA3-2xHA) or co-transfected with zebrafish Tlr4ba, Tlr4bb (a generous gift from Dr. Victoriano Mulero, Spain) (Sepulcre et al., 2009), or a human TLR4 expression clone (Addgene, cat. #13018). Additionally, an empty vector or human MD-2 (OriGene, cat. #RC204686) were transiently expressed. Transfections were done following the manufacturer’s protocol using jetPRIME reagent (Polyplus, cat. #CA89129-924) with half the indicated amount of DNA. The media in each well was replaced with fresh media 4 hours post-transfection. 48 hours post-transfection, media was aspirated, and the cells were treated with nickel (II) chloride hexahydrate (Sigma, cat. #379840), or LPS (Invitrogen, cat. #L23351) diluted in fresh media.

### 2.8 Western blot analysis of zebrafish Tlr4 expression

Transfected cells were placed on ice for 5-10 minutes, washed with ice-cold PBS, and returned to ice. Cells were lysed using 200 μL of cold Modified Oncogene Science Lysis Buffer [MOSLB; 10 mM HEPES at pH 7.4, 50 mM Na Pyrophosphate, 50 mM NaF, 50 mM NaCl, 5 mM EDTA, 5 mM EGTA, 100 μM Na_3_VO_4_, 1% Triton X-100, Roche protease inhibitor cocktail (Sigma, cat. #11697498001); kindly provided by the Marchant group, UofA MMI) and by rocking for 15-20 minutes. The wells were rinsed with the lysis buffer and the lysates were collected in pre-chilled microcentrifuge tubes. The lysates were centrifuged at 4ᵒC for 10 minutes at 6000g and then the supernatants were collected in new pre-chilled tubes. Lysates were mixed with 6X Laemmli buffer at a 1:5 (buffer:lysate) ratio and then the samples were loaded without heating and separated on a 10% SDS-PAGE gel before transfer to a nitrocellulose membrane. The membrane was blocked with LiCor Intercept (TBS) Blocking Buffer for 1 hour at room temperature, then probed with mouse anti-V5 antibody (1:5000) (Invitrogen, cat. #R96025) overnight at 4ᵒC, and then probed with goat anti-mouse secondary antibody (1:5000) (LiCor, IRDye 800CW) for 1 hour at room temperature. The membrane was washed with TBST and imaged on a LiCor Odyssey.

### 2.9 HEK293T cell viability and IL-8 secretion

IL-8 cytokine secretion relative to no agonist treatment was used as a measure of TLR4 in these cells. Supernatants were collected 24 hours post-treatment and then IL-8 secretion was quantified using commercial human IL-8 ELISA kits (Invitrogen, cat. #88-8086) according to the manufacturer’s protocol. IL-8 secretion was normalized to cell viability to account for cell death by nickel. Cell viability was measured using MTT reagent (3-(4,5-dimethylthiazol-2-yl)-2,5-diphenyl tetrazolium bromide) (ACROS, cat. #158990010). MTT was diluted to 1 mg/mL in fresh media and added to the cells post-treatment and incubated for 4 hours. Formazan was solubilized in DMSO (dimethyl sulfoxide) (Sigma, cat. #276855) and then the absorbance was measured at 590 nm on a SpectraMAX i3x plate reader (Molecular Devices).

### 2.10 Phylogenetic and synteny analysis of zebrafish Tlr4

A phylogenetic reconstruction analysis of TLR4 was performed in MEGA11 as described (Hall et al., (2013; Tamura et al., 2021). Briefly, multiple protein sequences from different vertebrates with e-values <0.001 were collected from NCBI through a BLASTp of zebrafish Tlr4ba in MEGA11. A multiple sequence alignment was performed using MUSCLE under the default options. To determine the most appropriate model for sequence evolution under maximum likelihood (ML) for this data set, we used MEGA’s program to find the best model for estimating the tree. The Nearest-Neighbor-interchange (NNI) algorithm was used to generate initial unrooted tree(s) for the heuristic search, which was estimated using the JJT+F +G +I substitution model with 1000 bootstrap replications (Jones et al., 1992). All positions with less than 95% site coverage were eliminated, i.e., fewer than 5% alignment gaps, missing data, and ambiguous bases were allowed at any position.

Synteny analysis was performed using the Genomicus 93.01 available at: https://www.genomicus.bio.ens.psl.eu/genomicus-93.01/cgi-bin/search.pl. The gene tree was rooted at the duplication of Tlr4 in early vertebrates. All Tlr4 sequences are collected from the Ensembl database, with all low-coverage sequences removed from the tree. Species for comparison were then selected for analysis.

### 2.11 Statistical analysis

Individual neuromast scores for each larva were added together to provide a neuromast viability score for each individual animal, and this score was used for subsequent analysis. The mean neuromast viability scores and hair cell counts from each group were analyzed using one-way ANOVA with Tukey’s multiple comparisons test. The EC_50_ for metal ion data was calculated by normalizing each data set, followed by a non-linear regression with a variable slope. When the data was normalized, 0% and 100% of a response was defined as the smallest and largest mean in each data set respectively. All graphs and statistical tests were performed using GraphPad Prism 9. HEK293T normalized IL-8 secretion values were analyzed using an RM two-way ANOVA and Tukey’s multiple comparisons test.

## 3. Results

### 3.1 Nickel, cobalt, and platinum induce hair cell death in larval zebrafish neuromast hair cells

In mammals, nickel is known to induce allergic contact hypersensitivity reactions through the direct binding and activation of TLR4 (Peana et al., 2017; Schmidt et al., 2010). Other group X transition metals, such as platinum, have recently been shown to activate mammalian TLR4 through a similar mechanism as nickel (Domingo et al., 2023). Previous findings show Tlr4 signalling is highly conserved between mammals and zebrafish, with the lowest conservation area being within the extracellular domain (Sullivan et al., 2009; Zhang et al., 2014) (Supplementary Figure 2). This difference, along with lack of co-receptors, is believed to contribute to the low sensitivity of zebrafish Tlr4 homologs to LPS. Instead, we speculate zebrafish Tlr4 might be suitable for the direct binding of metal ions. Indeed, we recently established that a platinum-based chemical is toxic to zebrafish neuromast hair cells in a manner that requires zebrafish TRL4 homologs (Babolmorad et al., 2021), so we reasoned that hair cell health could be a favourable proxy of zebrafish Tlr4 mediating metal toxicity *in vivo*. We examined the ototoxicity of NiCl_2_, CoCl_2_, PtCl_2_, and PtCl_4_ and found a monophasic dose-dependent relationship between neuromast viability and metal ion concentration (Figure 2). The EC_50_ of neuromast viability for NiCl_2_, CoCl_2_, PtCl_2_, and PtCl_4_ was found to be 7.6μM, 4.6μM, 0.44μM, and 0.64μM respectively, demonstrating the high toxicity of platinum salts to hair cells (Supplementary Figure 3). PtCl_2_ had a >10-fold toxicity to the pLL neuromasts in comparison to NiCl_2_, which was the least toxic metal compound (Figure 2D & 2F). Group IX/X transition metals did not appear to have any overt morphological effects on larvae over the course of treatment (applied for 20 hours at 5-6 dpf; Supplementary Figure 4). Together, these results demonstrate that the tested group IX/X transition metals induce dose-dependent neuromast hair cell death in larval zebrafish.

### 3.2 Zebrafish TLR4 homologs are required for metal toxicity in PLL neuromast hair cells

To test whether the cytotoxic effects on neuromast hair cells from nickel and platinum require zebrafish Tlr4, we used CRISPR-Cas9 gene editing technology to produce crispant larvae where all three zebrafish *tlr4* genes (*tlr4ba, tlr4al,* and *tlr4bb*) were targeted for mutagenesis. Crispant larva were established by injecting a Cas9/gRNA ribonuclear complexes (RNPs) targeting a sequence early in *tlr4ba*, *tlr4al*, and *tlr4bb* gene homologs (Supplementary Figure 5). The “mock” injected larvae received Cas9 protein but not the gRNA duplex and were used as a control for any effects of the microinjection process. CRISPR efficacy was confirmed by sequencing the *tlr4* genes. As expected, Sanger sequencing showed no detectable mutations within the *tlr4* genes of uninjected or mock injected larvae, while crispant larvae had non-consensus sequences beginning in their *tlr4ba* and *tlr4bb* genes at positions that exactly aligned with CRISPR gRNA binding sites, indicating genome cleavage and variable mutations products (Supplementary Figures 6-8). This confirmed that the *tlr4ba* and *tlr4bb* genes were successfully disrupted in crispants, whereas the status of *tlr4al* remains ambiguous. Crispant larvae displayed no overt phenotypes compared to uninjected, or mock injected larva prior to application of metals (Supplemental Figure 3).

We used these tlr4 crispants to test the hypothesis that zebrafish TLR4 homologs are required for mediating the metal toxicity observed in hair cells. Knockdown of TLR4 homologs protected hair cells from metal toxicity (Figure 3). Treatment with 10µM NiCl_2_, 0.75µM PtCl_2_ or 0.75µM PtCl_4_, led to significantly less damage to neuromast hair cells in crispants compared to controls (Figure 3). The mutations in *tlr4ba* and *tlr4bb* resulted in a greater than 2-fold increase in mean neuromast viability after treatment with nickel and platinum compounds in comparison to uninjected larva, demonstrating the involvement of zebrafish Tlr4 in metal induced neuromast hair cell toxicity.

**Figure 3.**
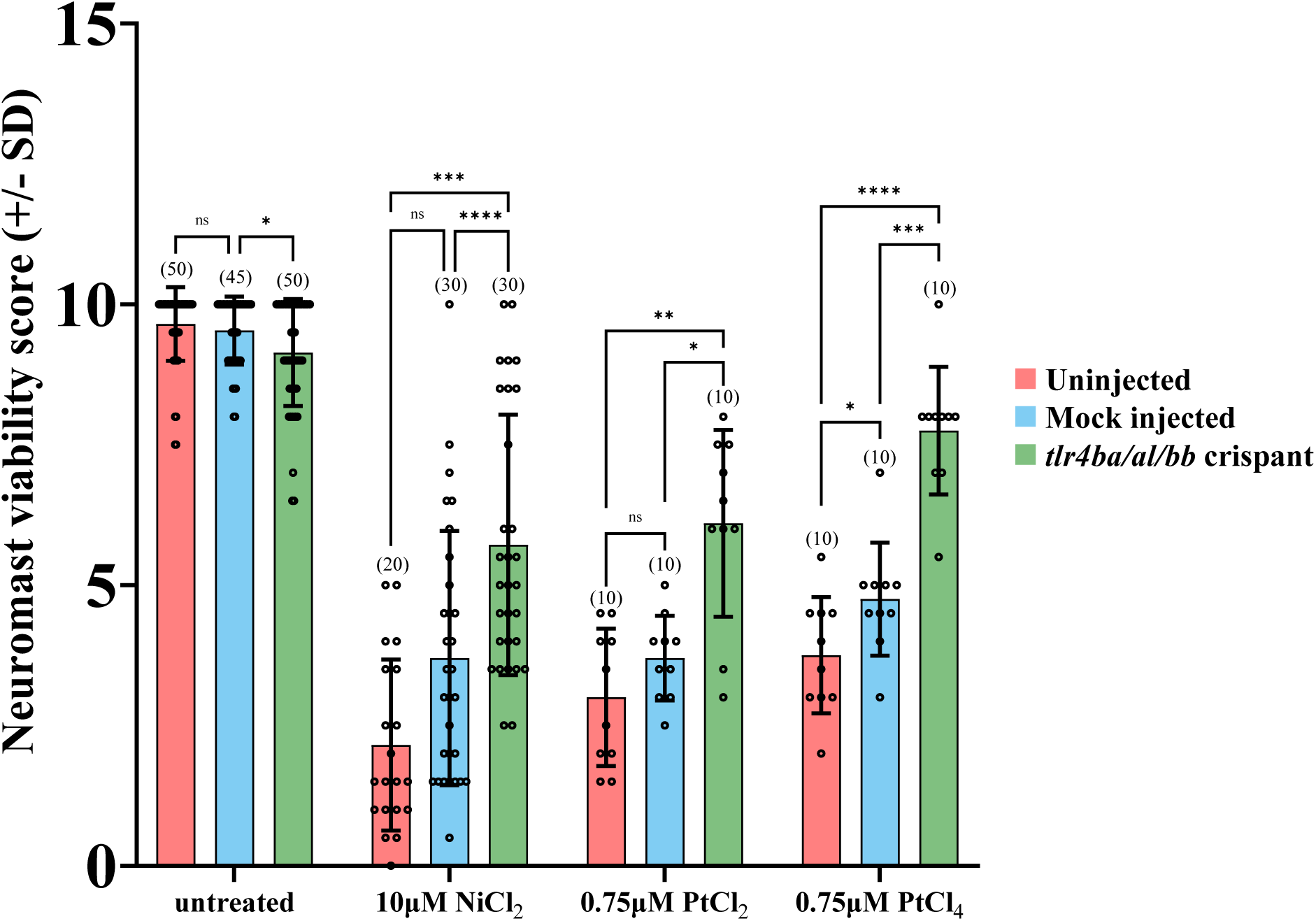
*tlr4* crispant mutant larvae are less susceptible to Group 10 transition metal induced ototoxicity. Both uninjected and mock injected larvae show a significant decrease in neuromast hair cell survival after treatment with 10μM NiCl_2_ and 0.75μM PtCl_2_ and PtCl_4_. The *tlr4* crispant larvae had their *tlr4* homologs mutated, protecting them from group 10 transition metal induced ototoxicity. Neuromast viability scores of tlr4 crispants were significantly higher than mock and uninjected nickel and platinum treated groups. Larvae were mutated in their *tlr4* homologs using CRISPR-cas9 genome editing. Larvae were injected at the single-cell stage with either gRNA sequences targeting the three zebrafish tlr4 homologs or Cas9 protein alone (mock). Larvae were treated with cisplatin for 20 hours at 6 dpf. Bracketed numbers above each bar represents the total number of larvae scored in each group. ****p<0.0001, ns = not significant by one way ANOVA and Tukey’s multiple comparison test.

### 3.3 Zebrafish Tlr4 transfected into HEK293T cells show a modest response to nickel, but no measurable response to LPS

HEK293T cells do not normally express TLR4 but can be rendered responsive to TLR4 agonists when transfected with human TLR4 and MD-2 – they have been shown to upregulate proinflammatory gene expression in response to known TLR4 agonists (Babolmorad et al., 2021; Chow et al., 1999; Domingo et al., 2023; Medvedev & Vogel, 2003; Yang et al., 2000). IL-8 has been established by numerous studies as a marker for TLR4 activation making HEK293T cells an appropriate *in vitro* cell model to assess TLR4 function (Kurt-Jones et al., 2000; McKee et al., 2021; Oblak et al., 2015; Potnis et al., 2013; Quevedo-Diaz et al., 2010; Schmidt et al., 2010).

Transient expression of human *TLR4* in our HEK cell-based system conferred responsiveness to nickel and LPS, in an MD-2 dependent manner, serving as a positive control for TLR4 activation in this system (Figure 4A). When similarly expressed, both zebrafish TLR4 homologs *tlr4ba* or *tlr4bb* were sufficient to mediate signalling to nickel (Figure 4B, C), approximately doubling the response. Stable expression of zebrafish homologs in the HEK cell system was confirmed by immunoblot (Supplementary Figure 9). Zebrafish homologs of TLR4 showed no measurable response to LPS (despite co-expression of human *MD-2*), consistent with past reports (Sepulcre et al., 2009; Sullivan et al., 2009).

**Figure 4.**
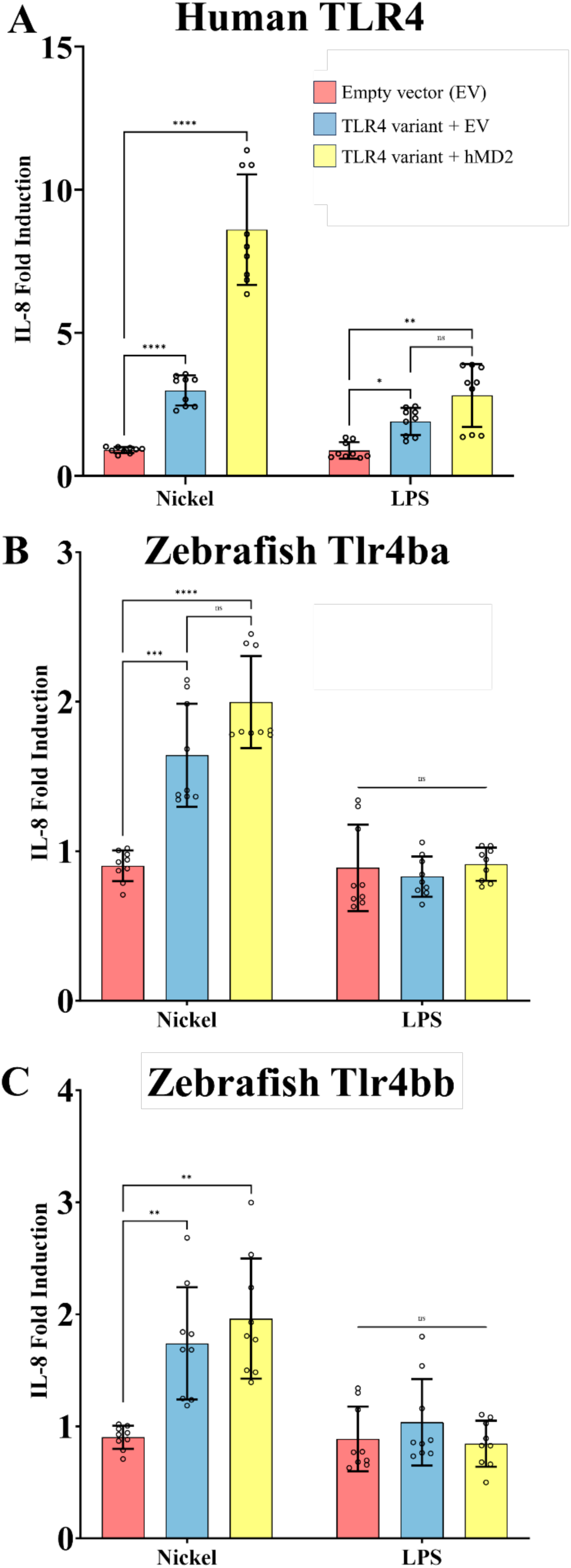
Nickel activates zebrafish Tlr4 homologs when expressed in a heterologous system. HEK293T cells transiently expressing *TLR4* homologs from human (panel A) and zebrafish (Tlr4ba and Tlr4bb, in panels B and C, respectively) with and without MD-2 were treated with the potential TLR4 agonists nickel (200 µM) or LPS (10 ng/mL). (**A**) Human TLR4 confers responsiveness to Ni or LPS agonists, as measured via IL-8 secretion, serving as a benchmark comparator for zebrafish homologs. (**B,C**) Zebrafish Tlr4ba and Tlr4bb confer responsiveness to nickel as assessed by IL-8 secretion, but are largely independent of MD-2 expression. Data are presented as the fold increase relative to untreated isogenic samples. N = three independent experiments for all treatments. ****p<0.0001, ***p<0.0002, **p<0.0021, *p<0.0332, ns = not significant by two-way RM ANOVA and Šídák’s multiple comparisons test.

## 4. Discussion

This study demonstrates that group IX/X transition metals mediate the activation of zebrafish homologs of TLR4, providing insight to the long-standing mystery of TLR4 function in early-branching vertebrates. We showed that bath application of NiCl_2_, CoCl_2_, PtCl_2_, or PtCl_4_ impacted the viability of neuromast hair cells and that this effect required zebrafish homologs of TLR4. Furthermore, zebrafish Tlr4ba or Tlr4bb were capable of transducing signals following nickel treatment when expressed in a heterologous cell platform. Altogether, this data suggests zebrafish homologs of TLR4 mediate cellular responses to metal ions, which is a response previously observed only in mammals (including humans and closely related primates).

Exposure to trace metals such as nickel, cobalt, zinc, and copper has been previously observed to induce inflammatory responses in zebrafish (Juśkiewicz & Gierszewski, 2022; Singh et al., 2023). Metal ions induce expression of proinflammatory cytokines such as *il1β* and *tnfα* as well as Tlr signalling genes *myd88*, *nfkb1a*, and *tlr5b* (Brun et al., 2018; Chen et al., 2019; Wang et al., 2015). The involvement of Tlrs in mediating these inflammatory responses is further supported by the observation that the MAPK pathway also shows increased gene expression after metal exposure (Chen et al., 2019). In humans, metals elicit these responses described above through the binding and activation of TLR4 (Babolmorad et al., 2021; Oblak et al., 2015; Rachmawati et al., 2013; Schmidt et al., 2010). Additionally, as observed in humans following metal-induced TLR4 activation, contact with metal ions in zebrafish can lead to the development of high concentrations of reactive oxygen species (ROS), leading to further inflammation, disruption of mitchondrial function, and eventually cell death (Jia et al., 2022; Wang et al., 2015). In this study we show that during bath application with metal ions, clusters of hair cells on the exterior of zebrafish, called neuromasts, aptly demonstrate the ability of metals to induce cell death. This effect can be mitigated by mutations in Tlr4ba/Tlr4bb, resulting in a reduction of metal cytotoxicity. This data provides evidence that the generation of metal-induced cell death in zebrafish is mediated in some part through the activity of Tlr4 homologs.

These findings provide further support to our hypothesis that zebrafish Tlr4 can be activated by metals, and while this suggests an ancient, shared function for vertebrate TLR4 it also raises new questions about how previous works align with this conclusion. The binding of nickel to human TLR4 has been shown to be directly mediated through two histidine residues within the LRR domain. These residues are conserved between primate species such as chimpanzees and humans but are largely absent in more distantly related vertebrates (Peana et al., 2017; Schmidt et al., 2010). Previously, it has been suggested that due to the lack of these histidine residues, other species apart from primates fail to respond to metal ions through TLR4 (Oblak et al., 2015; Schmidt et al., 2010). Mice have demonstrated nickel hypersensitivity through TLR4 activation, but only when pre-sensitized with LPS as an adjuvant (Sato et al., 2007). In contrast, our findings show that cells transfected with zebrafish Tlr4 homologs increase secretion of IL-8 in response to nickel and in the absence of LPS, similar to what is seen in humans (Rachmawati et al., 2013; Schmidt et al., 2010). Furthermore, MD-2 was not required for metal induced activation of zebrafish Tlr4, suggesting a binding mechanism like that observed in humans. Further work is warranted towards understanding which residues in zebrafish Tlr4 mediate metal signalling. Regardless, this implicates zebrafish as a promising animal model for studying metal allergies and provides an in vivo system to examine the complexity of metal-induced contact hypersensitivity reactions and future therapeutic avenues.

Human TLR4 is a promiscuous receptor known to be activated by a variety of ligands beyond LPS and nickel, including one of its most important functions: recognition of cellular damage following the release of damage-associated molecular patterns (DAMPs). Metal ions may result in the release of DAMPs into the media, which then mediate the activation of zebrafish Tlr4 homologs. DAMP release through the generation of ROS (either through TLR4 or directly by metals) will result in positive feedback loop where activation of TLR4 leads to oxidative stress, leading to the release of DAMPS, which further activates TLR4. Therefore, we cannot rule out the possibility of DAMPs as a confound within this study. However, the activation of hTLR4 by some DAMPS such as HMGB1, requires co-receptors such as MD-2, which is not required for IL-8 secretion by cells expressing zebrafish Tlr4 (Yang et al., 2015). Furthermore, to our knowledge zebrafish Tlr4 homologs have only one endogenous ligand, but examining endogenous activators of zebrafish Tlr4 provides an exciting avenue for future study. For example, Liu et al. showed that peroxiredoxin 1, a widely expressed antioxidant enzyme and DAMP, could interact with Tlr4ba, inducing NF-κB activity and upregulating expression of proinflammatory cytokines (He et al., 2019; 2018). This finding supports some role of DAMPs in zebrafish Tlr4 activation, however more work is required to determine the role DAMPs play in metal induced Tlr4 activation in zebrafish.

The ability of zebrafish Tlr4, particularly Tlr4ba, to respond to LPS was recently re-visited by Loes et al. after their insightful observation of an MD-2 homolog present within the zebrafish genome (2021). LPS was shown to be capable of activating zebrafish Tlr4ba in the presence of MD-2, however, there are two caveats: it required mammalian Cd14, which currently has no known homolog within zebrafish, and the concentrations required for activation of zebrafish Tlr4 were considerably higher (μg/mL) than the biologically relevant concentrations of LPS (0-10ng/mL) required for TLR4 activation (Guo et al., 2013). Importantly, LPS has complex variations between sources and preparations; Further exploration is needed to determine if LPS sourced from fish pathogens might be better able to initiate signalling of fish TLR4 homologs.

On the other hand, dimerization by metal ions is through a direct binding mechanism and excludes the aforementioned prerequisite of CD14 for activation. The presence of Tlr4 homologs and an MD-2 homolog within zebrafish, along with LPS sensitivity being undetectable, provided the logical basis for our hypothesis that zebrafish Tlr4 utilizes group IX/X metals for ligands. The most recent hypotheses for how zebrafish Tlr4 fits in to the evolution of this receptor suggests the *tlr4* gene along with *ly96* (MD-2) arose in a common ancestor before divergence of lobe-finned and ray-finned fish (Loes et al., 2021). The lack of a CD14 homolog, and the ill-defined LPS sensitivity in most fish species, further supports the notion that Tlr4’s purpose originally included the binding of ligands that do not require co-receptors, such as metals.

In summary, we present data in support of nickel and platinum acting as novel ligands for zebrafish Tlr4, helping discern the mystery surrounding the function of TLR4 across evolutionary time. We demonstrate that genetic mutation of Tlr4 renders zebrafish significantly less sensitive to metals, whereas zebrafish homologs of TLR4 when expressed in heterologous cells were sufficient to induce an inflammatory response to metals. These findings suggest a previously unknown function of zebrafish Tlr4, which includes detecting metals in their environment or physiology. We anticipate that an improved understanding of the function of zebrafish Tlr4 and its involvement with metal ions will facilitate better utilization of this model organism in the study of innate immunity and the myriad of various diseases involving inflammation, particularly including metal contact hypersensitivity.

## Acknowledgements

Victoriano Mulero kindly provided plasmids for expression of zebrafish homologs of zebrafish TLR4. Operating grants to APB and WTA from CIHR (PJT-178327). Studentship to TSL from the Cancer Research Institute of Northern Alberta. We appreciate support of animal wellbeing provided by Science Animal Support Services at the University of Alberta. Chris Li assisted with generation of gRNA sequences as well as PCR primers for crispant validation.

## Supplemental Materials

**Supplementary Figure 1.**
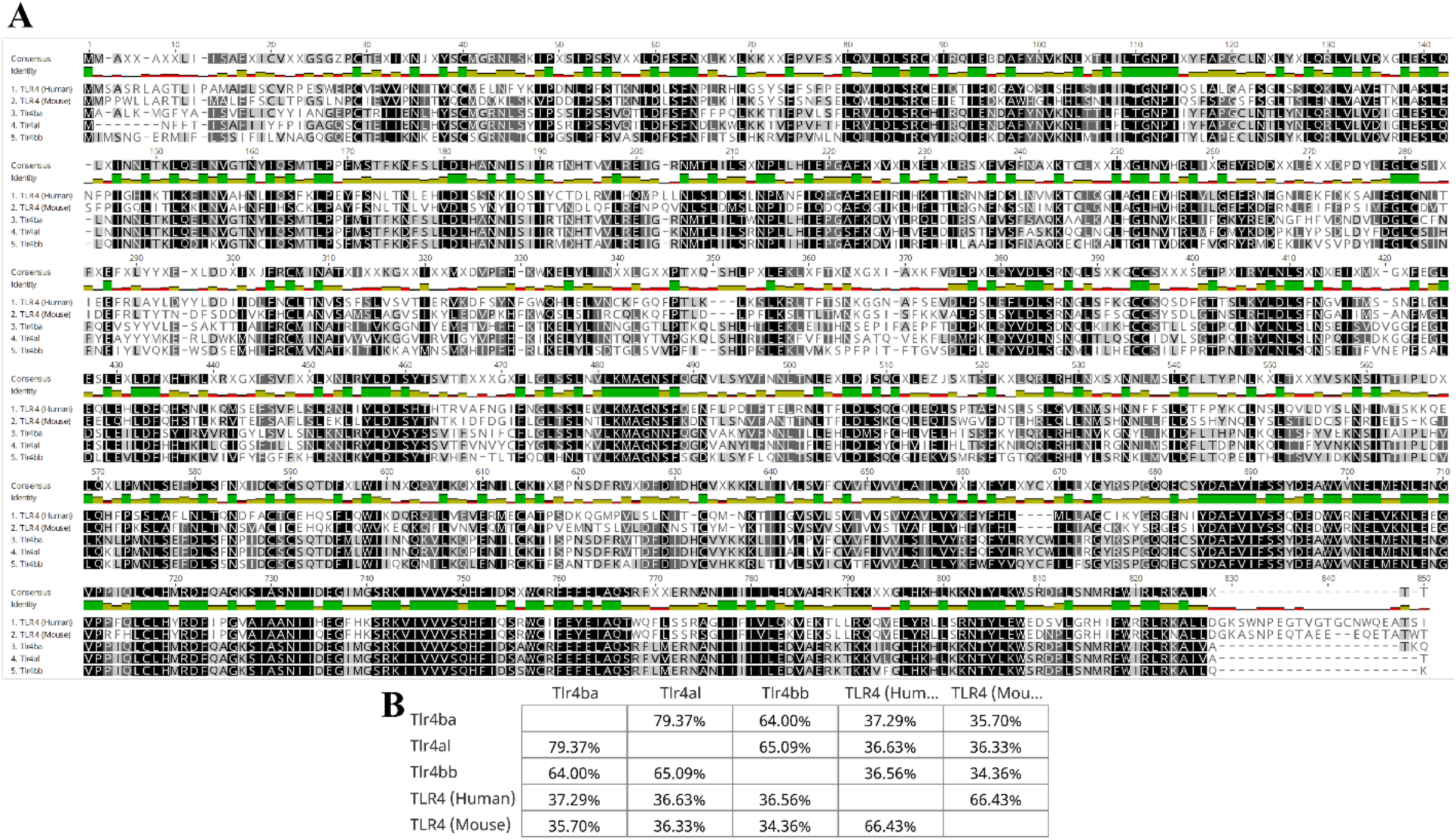
Multiple sequence alignment and percent identity matrix for TLR4 (full protein) between zebrafish, mice, and humans. A) The comparison between sequences shows the homology and conservation of residues mainly in the latter third of the protein sequence. Similar residues (100%) are highlighted in black, partially similar residues are highlighted in dark grey (80%-100%) and light grey (60%-80%). The coloured histogram shows the mean pairwise percent identity for each column of the alignment, where green represents 100%, yellow represents >30%, and red represents <30%. B) Table shows the percent sequence identity between mammalian TLR4 (mouse and human) is >60%, while the percent identity between mammalian and zebrafish Tlr4 homologs is <40%. Alignments, similarity, and the percent identity matrix were made and calculated using Geneious Prime.

**Supplementary Figure 2.**
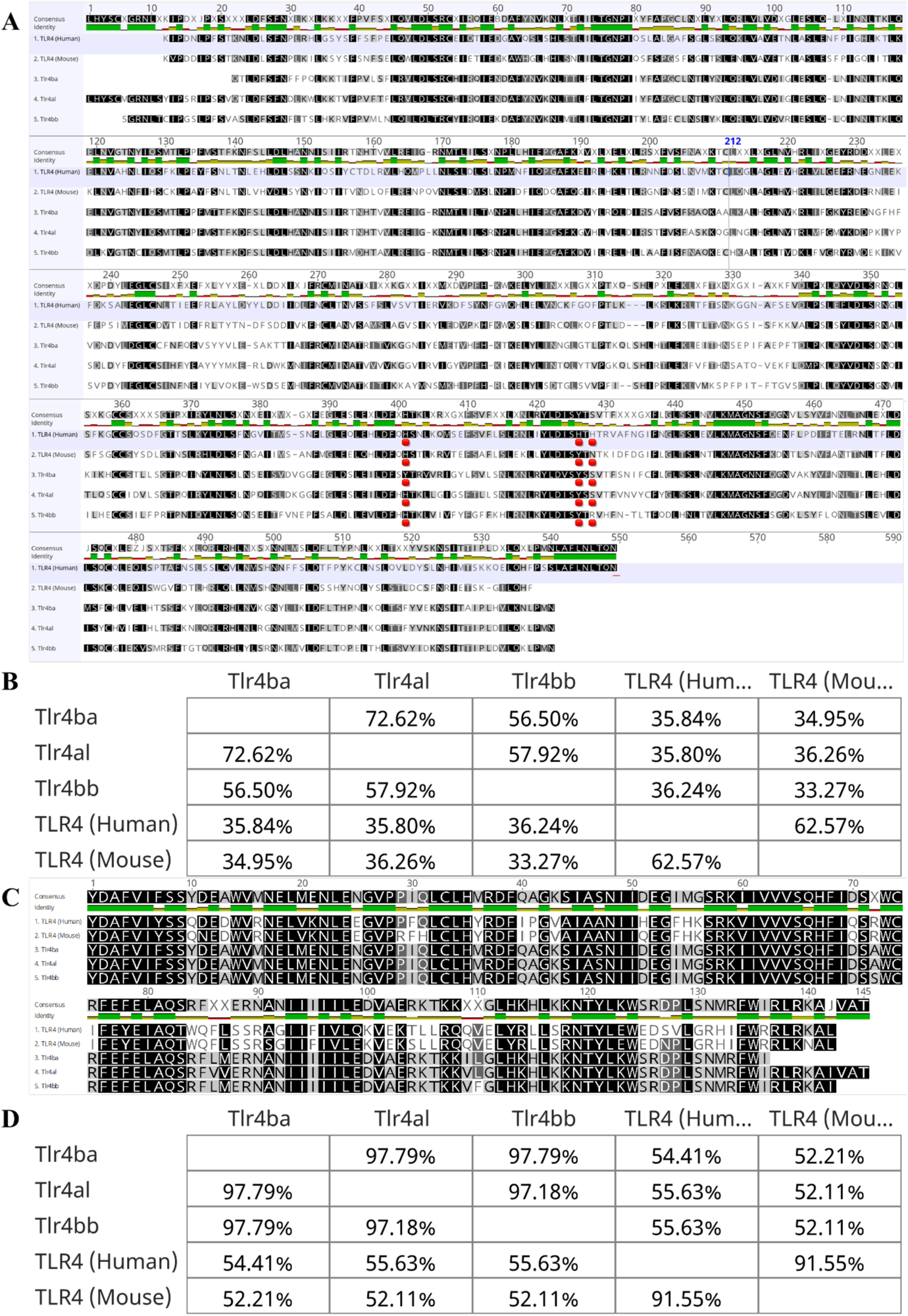
Multiple sequence alignment and percent identity matrix for TLR4 extracellular LRR domain and intracellular TIR domain between human, mouse, and zebrafish. The comparison between extracellular LRR domain (A, B) sequences shows the homology and conservation of residues is ∼20% greater within the intracellular TIR domain (C, D) compared with the extracellular LRR domain. Similar residues (100%) are highlighted in black, partially similar residues are highlighted in dark grey (80%-100%) and light grey (60%-80%). The coloured histogram shows the mean pairwise percent identity for each column of the alignment, where green represents 100%, yellow represents >30%, and red represents <30%. The known human nickel binding residues are underlined in red within A. The nickel binding residues 431 and 456 show 100% similarity between humans, mice, and zebrafish, as well as partial identity (>30%) and consist of either histidine or tyrosine residues. The last known nickel binding residue (458) shows low similarity and identity between all sequence and is either a histidine, asparagine, serine, or arginine residue. The tables (B, D) below the alignments (A, C) show the percent sequence identity for the entire sequence alignment between human TLR4, mouse TLR4, zebrafish Tlr4ba, Tlr4al, and Tlr4bb. Alignments, similarity, and the percent identity matrix were made and calculated using Geneious Prime.

**Supplementary Figure 3.**
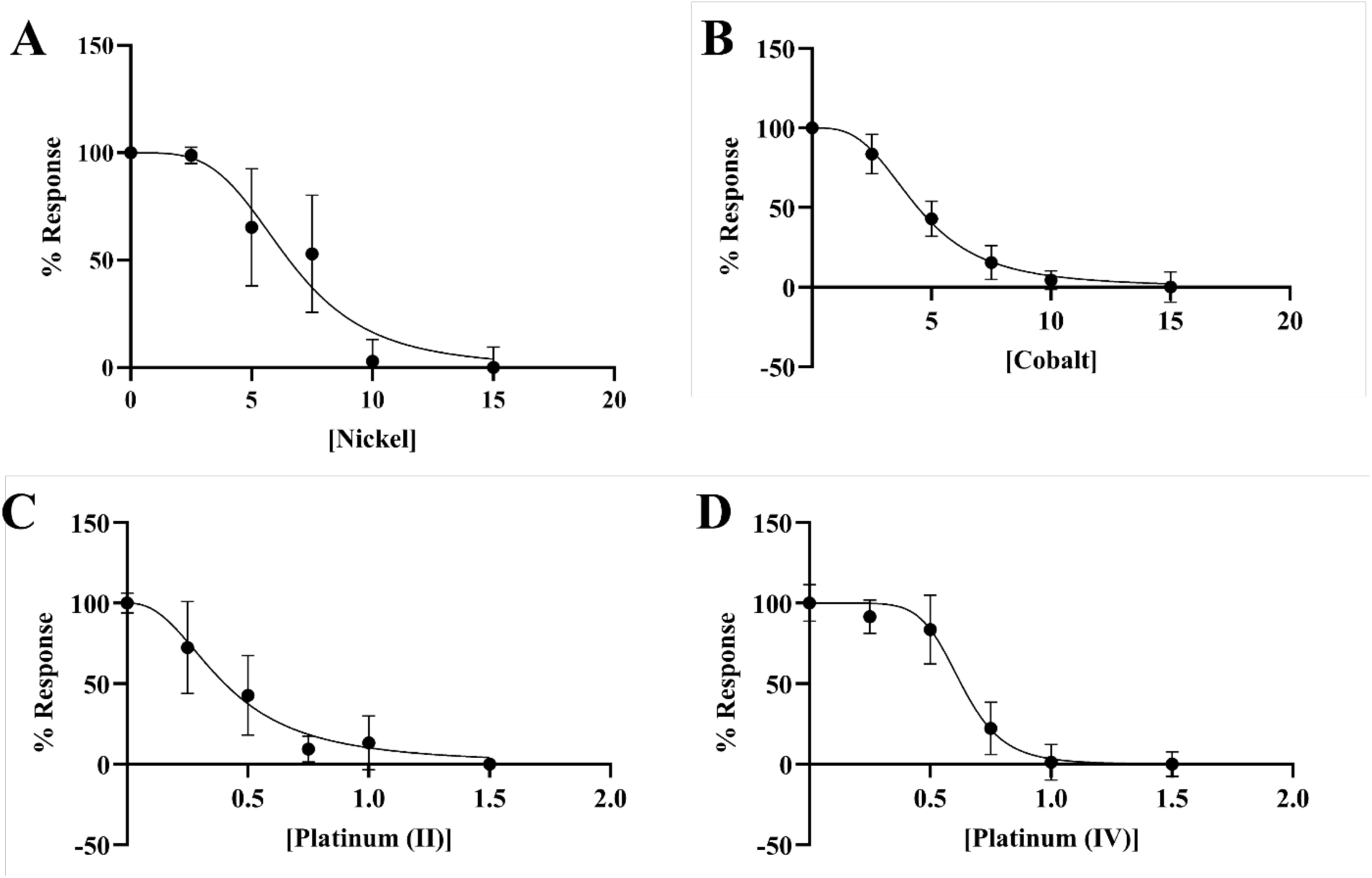
Normalized linear regression plots of nickel, cobalt, platinum II, and platinum IV dose response curves. Neuromast viability responses to varying concentrations of NiCl_2_ (A), CoCl_2_ (B), PtCl_2_ (C), and PtCl_4_ (D) were averaged and normalized where 0% and 100% of a response was defined as the smallest and largest mean in each data set respectively. EC_50_ values were then calculated using a non-linear regression with variable slope in GraphPad Prism 10.

**Supplementary Figure 4.**
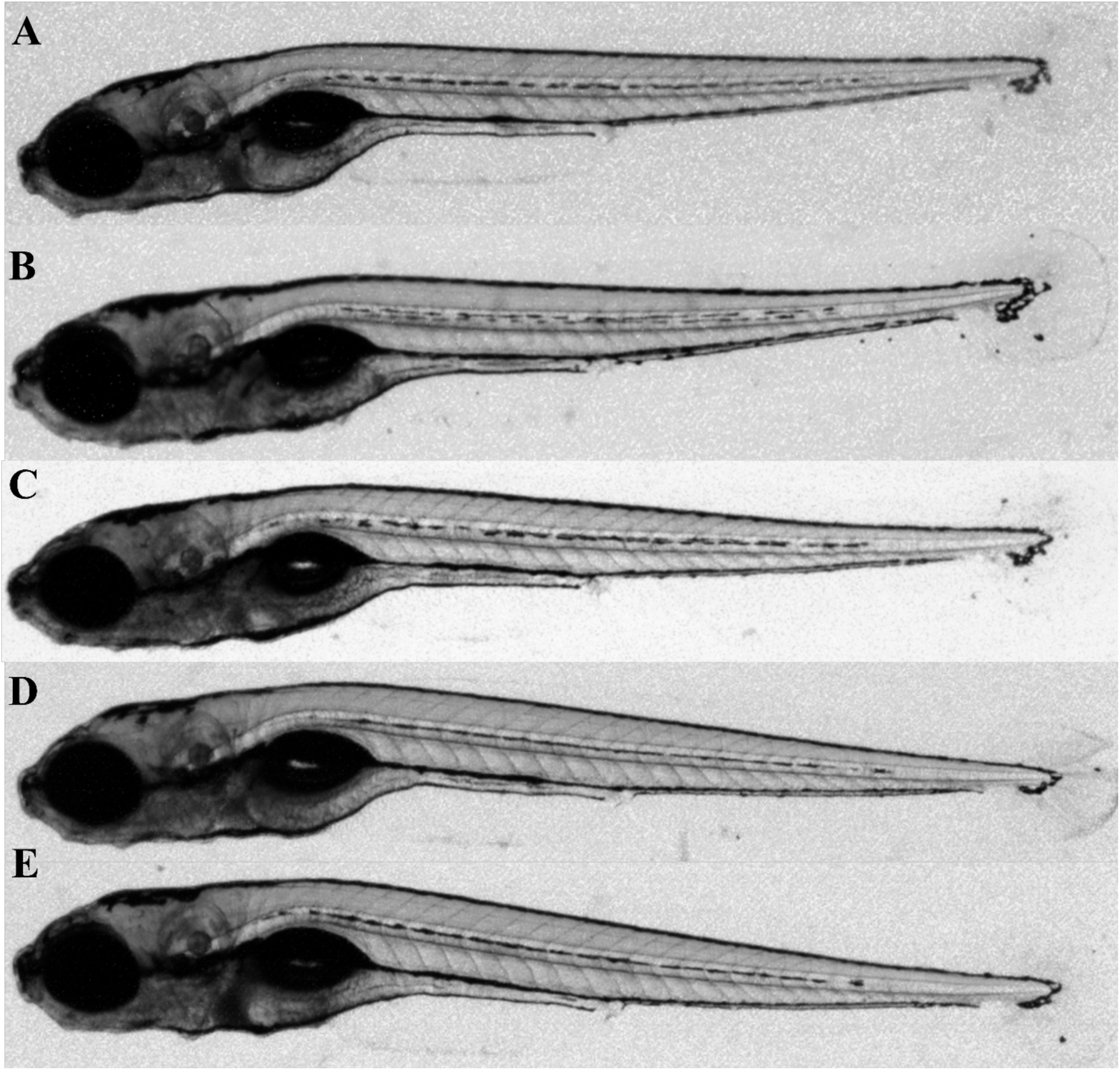
Tlr4 crispant fish treated with nickel (II) and platinum (IV). A-C) 6-7dpf mock injected and Tlr4 crispant larva treated with either PtCl_4_ or NiCl_2_ show no apparent morphological change in comparison to an uninjected untreated larva. A) Uninjected, untreated larva. B) Mock injected larva treated with 7.5µM PtCl_4_. C) Tlr4 crispant treated with 7.5µM PtCl_4_. D) Mock injected larva treated with 10µM NiCl_2_. E) Tlr4 crispant larva treated with 10µM NiCl_2_.

**Supplementary Figure 5.**
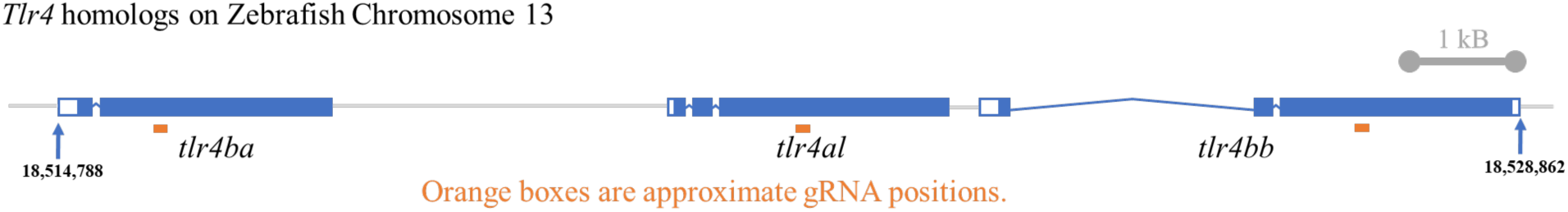
Illustration representing the three *TLR4* homologs on zebrafish chromosome 13 and their approximate gRNA positions. The three zebrafish *tlr4* homologs are found in succession to one another. Arrows point to the initial base pair position on the chromosome. White boxes represent untranslated regions, blue boxes represent exons, while blue lines denote introns. Grey lines represent the intergenic region connecting the homologs.

**Supplementary Figure 6.**
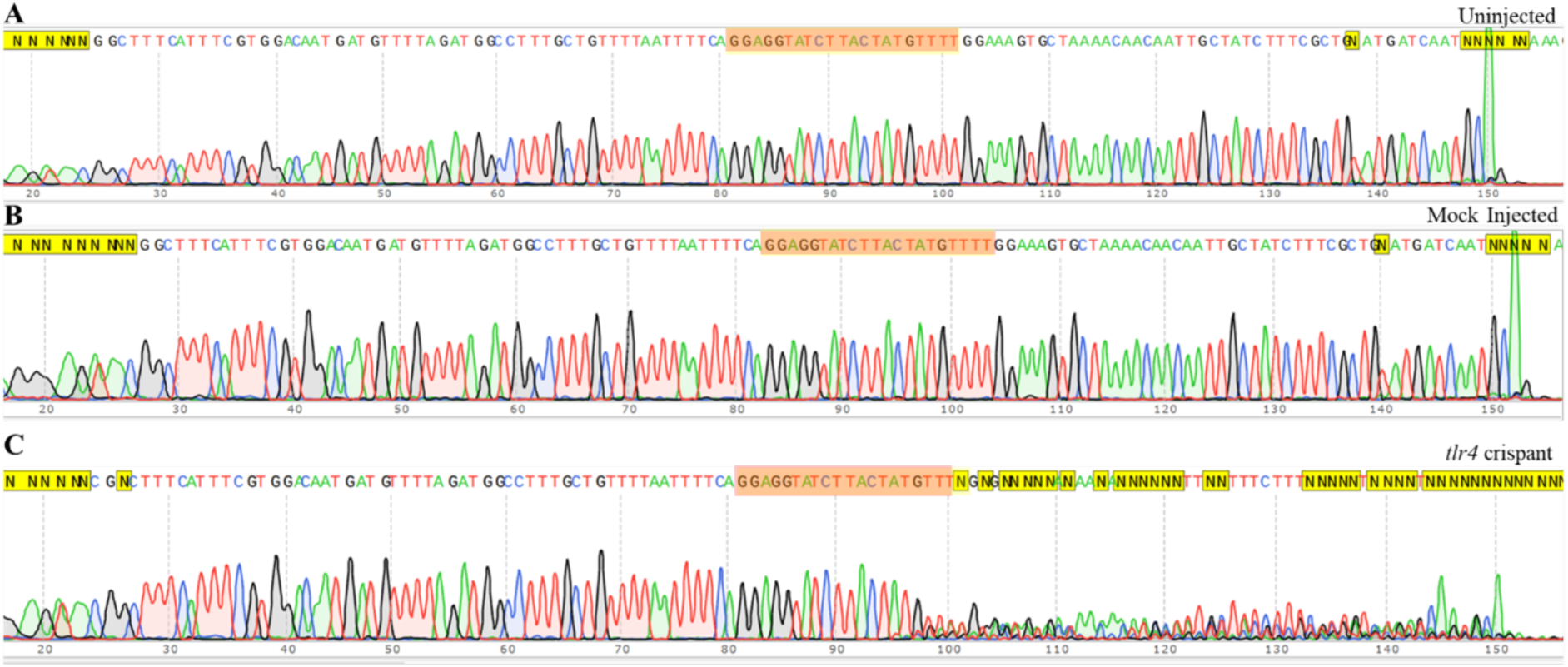
Sanger Sequencing of tlr4ba crispant DNA. The *tlr4ba* gene was amplified from the genomic DNA of individual larvae via PCR using the tlr4ba forward primer. Uninjected larva (A) and mock injected larva (B) DNA shows an unmutated *tlr4ba* sequence, where consistent DNA in all cells produces a coherent chromatogram throughout. C) Chromatograms from *tlr4* cripants show consensus sequence chromatograms where the DNA is homozygous and consistent throughout the individual (left side), but the chromatogram becomes discrepant from wild type sequence near the gRNA binding site (highlighted in red) because a mixture of various mutant DNA sequences now exists among the larva’s cells. Larvae were injected at the single-cell stage with either a gRNA sequence targeting the zebrafish tlr4ba homolog or Cas9 protein alone (mock).

**Supplementary Figure 7.**
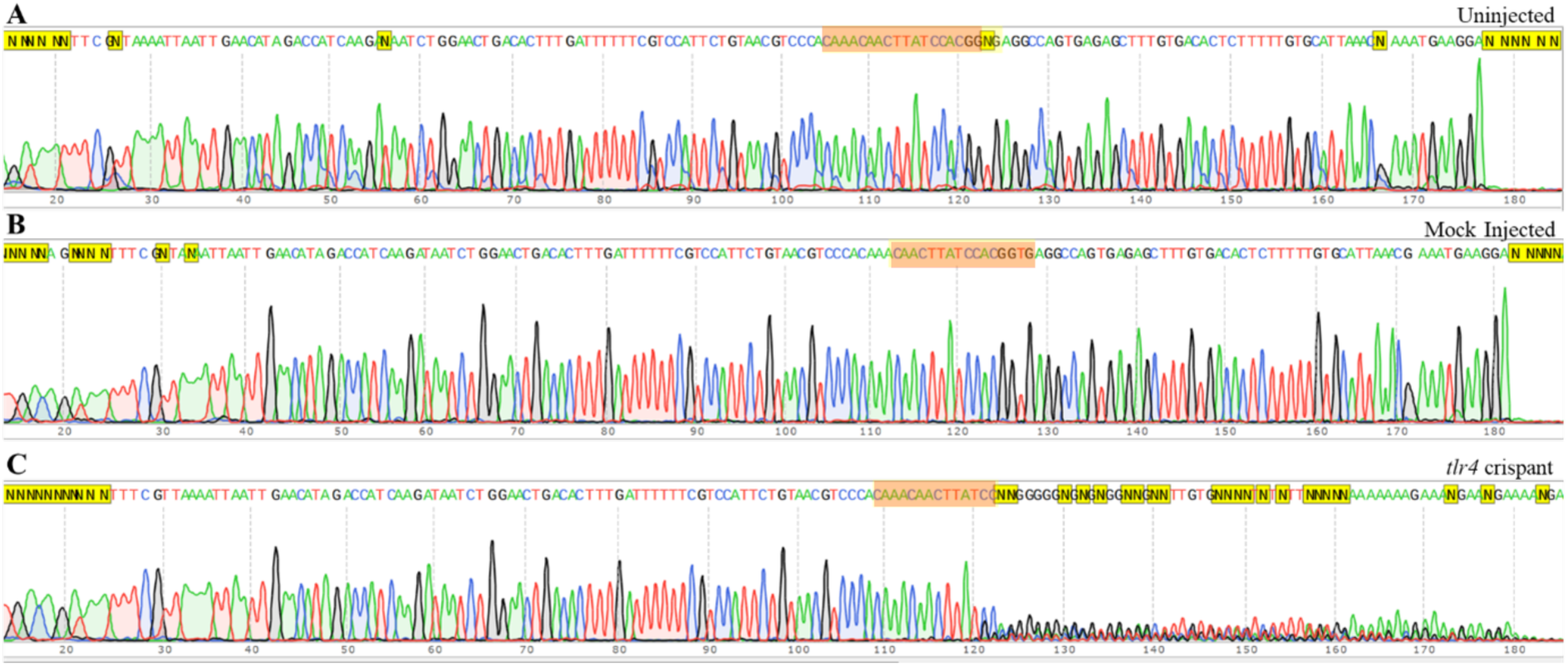
Sanger Sequencing of tlr4bb crispant DNA. The *tlr4bb* gene was amplified from the genomic DNA of individual larvae via PCR using the *tlr4bb* reverse primer. Uninjected larva (A) and mock injected larva (B) DNA shows an unmutated *tlr4bb* sequence, where consistent DNA in all cells produces a coherent chromatogram throughout. C) Chromatograms from *tlr4* crispants show consensus sequence chromatograms where the DNA is homozygous and consistent throughout the individual (left side), but the chromatogram becomes becomes discrepant from wild type sequence near the gRNA binding site (highlighted in red) because a mixture of various mutant DNA sequences now exists among the larva’s cells. Larvae were injected at the single-cell stage with either a gRNA sequence targeting the zebrafish tlr4bb homolog or Cas9 protein alone (mock).

**Supplementary Figure 8.**
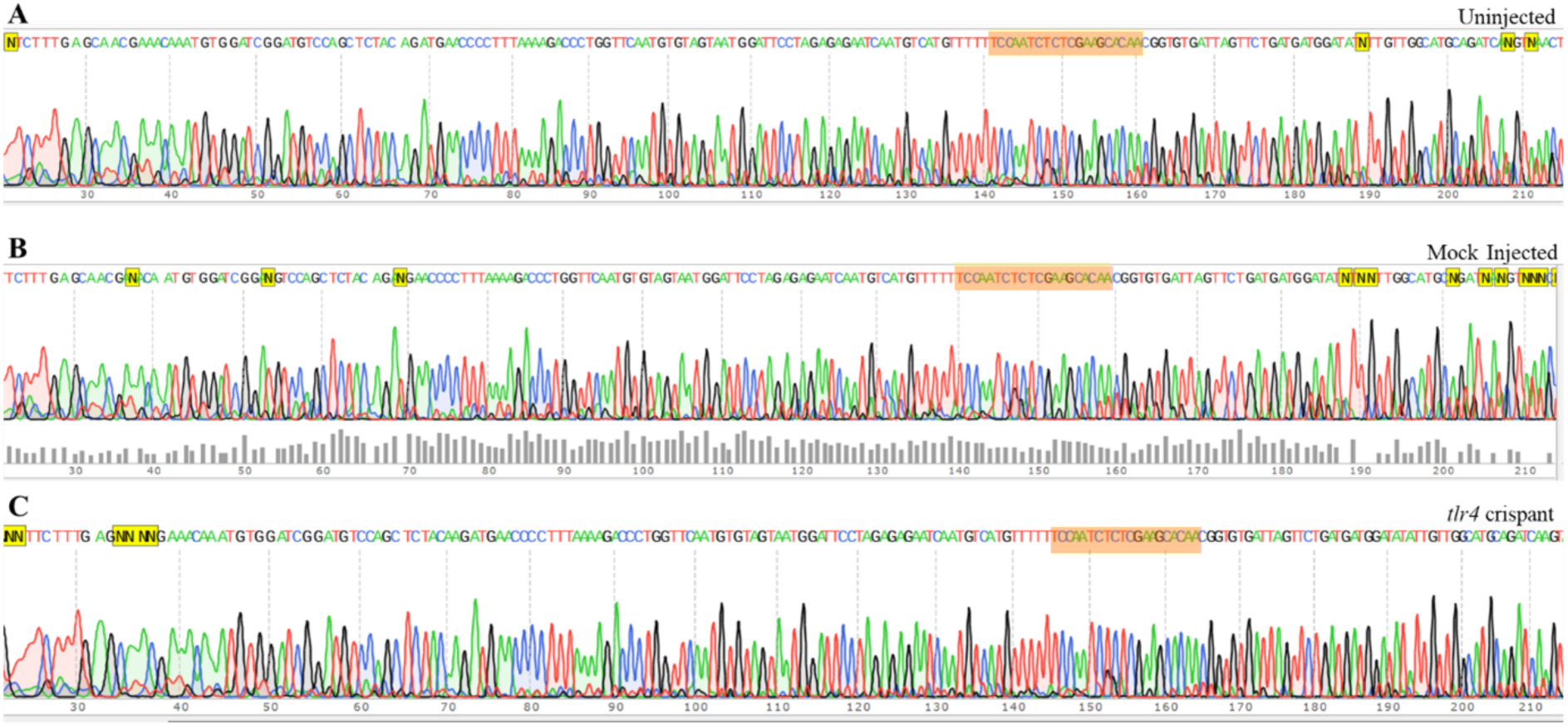
Sanger Sequencing of tlr4al crispant DNA. The *tlr4al* gene was amplified from the genomic DNA of individual larvae via PCR using the *tlr4al* reverse primer. Uninjected larva (A) and mock injected larva (B) DNA in A & B shows an unmutated *tlr4al* sequence, where consistent DNA in all larval cells produces a coherent chromatogram throughout. C) Chromatograms from *tlr4* cripants show consensus sequence chromatograms where the DNA is homozygous and consistent throughout the individual even following the gRNA target sequence (highlighted in red).

**Supplementary Figure 9.**
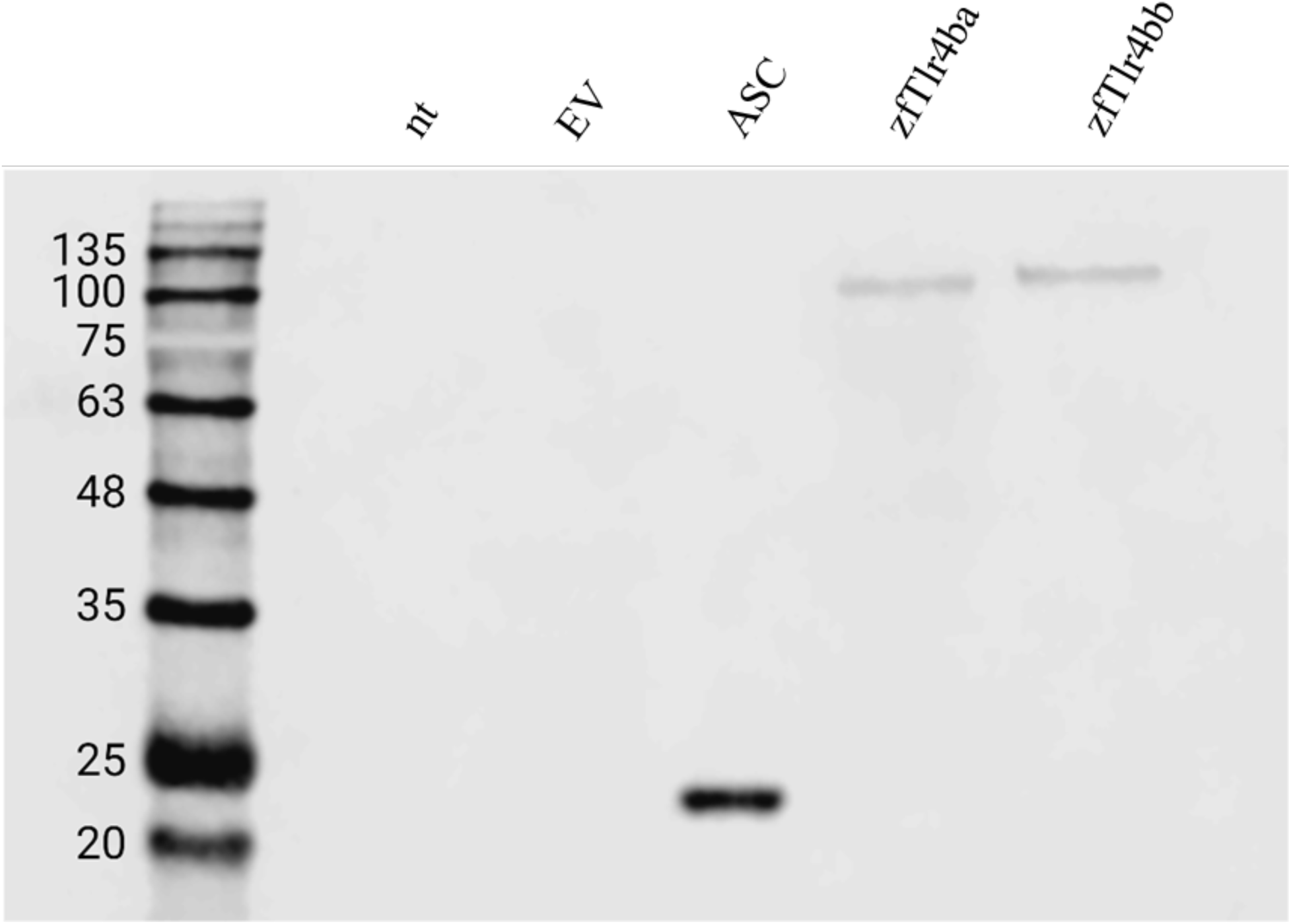
Zebrafish Tlr4ba and Tlr4bb are stably produced in HEK293T cells. Zebrafish *tlr4ba* and *tlr4bb* were transiently expressed in HEK293T cells from a constitutive promoter. A signal of the expected size was observed in cells transiently expressing V5/His_6_-epitope tagged zebrafish Tlr4 proteins (zfTlr4ba, zfTlr4bb). Lysate from cells transiently expressing V5-tagged ASC was used as a positive control, while lysates from non-transfected (nt) or empty vector (EV)-transfected HEK293T cells were used as negative controls.

**Supplementary Table 1.**
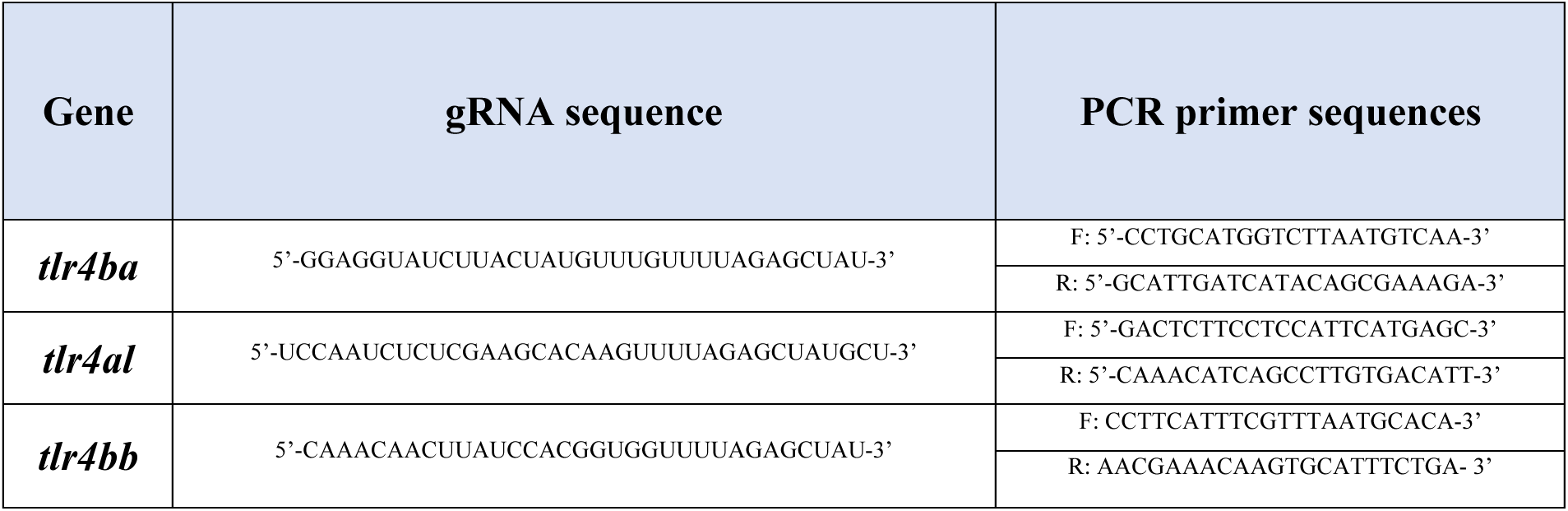
CRISPR-cas9 gRNA sequences and PCR primer sequences.

